# Making monkeys of caregivers: Evaluating the use of caregiver behaviour as social enrichment for lab-housed Rhesus and Cynomolgus Macaques

**DOI:** 10.64898/2026.11.16.767050

**Authors:** Rachel van Vliet, Nadege Aigueperse, Elsa Vasseur

**Affiliations:** Department of Animal Science, McGill University, Sainte-Anne-de-Bellevue, Quebec, Canada; INRAE, VetAgro Sup, UMR Herbivores, F-63122 Saint Genes-Champanelle, France

**Keywords:** Human-animal interactions, welfare, laboratory primates, social enrichment

## Abstract

Enrichment is pivotal to captive animal welfare. Social enrichment is of particular importance for social animals who, due to laboratory space or protocols, are not housed in larger groups. The objective of our study was to see if caregivers could act as an additional source of social enrichment to pair-housed research macaques. 5 adult Rhesus *(Macaca Mulatta)* and 3 adult Cynomolgus macaques *(Macaca fascicularis)* were enrolled in the study. Every day, for 3 weeks, caregivers would spend 6 minutes interacting with the monkeys during afternoon feeding. Half the monkeys received human behaviour (HB), where the caregivers interacted with the monkeys using human body language and speaking softly. The other half received monkey-like behaviour (MB), in which caregivers would attempt to imitate monkey behaviour, all the while avoiding eye-contact or speaking (both of which may be perceived as aggressive). Our results showed that monkeys were more likely to engage in interactions before feeding, indicating that this may be an ideal time to provide social enrichment. Monkeys from both groups showed a 3.5 ± 1.5 (F_54_ = 3.34, p = 0.058) decrease in the frequency of cage-directed abnormal behaviours and a 1.2 ± 0.53 (F_54_ = 2.78, p = 0.067) decrease in the frequency of stereotypic abnormal over the course of the experiment. In addition, all monkeys saw a 12.9 ± 6.1% (F_54_ = 2.40, p = 0.010) increase in the proportion of time spent interacting positively with cage-mates, including a 10.6 ± 4.78% (F_54.1_ = 2.47, p = 0.077) increase in time spent grooming. This research contributes to the enhancement of animal care by helping to establish evidence-based guidelines for the proper care of non-human primates in research. It further deepens our understanding of social enrichment provided by humans as a cost-effective way of managing boredom.

## 1. Introduction

Non-human primates are incredibly important research subjects in North America due to their close genetic relation to us and heightened cognitive capacities. While these traits make them invaluable for certain types of research (Servik 2018), they are also what make them particularly sensitive to the stressors of laboratory environments. Not only do non-human primates have higher cognitive capacities and social needs than many other research animals, making them more prone to boredom (Wemelsfelder 2005), but they have relatively long lifespans, subjecting them to aversive environments for longer periods of time, allowing negative effects to compound (CCAC 2019b). The most commonly used species of non-human primates in research are Cynomolgus macaques *(Macaca fascicularis)* and Rhesus macaques *(Macaca mulatta)* (Lankau et al. 2014).

Despite growing public demand for reduced use of primates in scientific research, non-human primates are seen as irreplaceable for certain models of cognitive behavior and human disease research (Servik 2018). This does not mean that steps have not been taken to improve captive environments. Rather, growing support for proper welfare accommodations of research animals from the research community, as well as outside voices, have pushed for improved conditions in research environments. In addition to animal ethics concerns, recent studies have revealed that subjects with a higher degree of welfare do in fact make for better science, as they represent healthy, normal individuals rather than psychologically and physically weakened ones (Buchanan-Smith 2010). A need therefore exists for increased studies on non-human primate welfare in order to propose better care and housing for non-human primates used in research.

One of the main methods used to mitigate the negative effects of captivity for non-human primates is environmental enrichment. This is a broad term which refers to any addition that can be made to a captive environment that improves that captive individual’s quality of life (UFAW n.d.). Social enrichment is of particular importance for non-human primates, since they are highly social animals. Unfortunately, the reality of research environments is that due to study protocols, animal health, or space availability, social housing is not possible and when it is, pair-housing is most common (Hannibal et al. 2017). This is a far cry from the complex social environments macaques experience in the wild, making it nearly impossible for research environments to adequately provide for macaques’ complex social needs. In order to accommodate for this, research has begun to look into using caregivers, who are already present in the daily lives of the captive primates, to provide additional social enrichment. The active use of caregivers as social enrichment is based on the idea of manipulating the relationships that naturally form between staff and captive primates (Claxton 2011).

The role of human caregivers in the lives of captive primates has changed significantly in the past 20 years, from protocols aiming to reduce human presence altogether, to an acknowledgement that they can play a positive role in the daily lives of the individuals in their care (Coleman 2011). The former more historic view is not without merit, however, and should still be taken into consideration. Humans, particularly unfamiliar or untrained ones, can and do present stressful stimuli to captive primates (Lambeth 1997, Bethell et a. 2012). In a review of the influence of humans in the refinement of non-human primate use in research, Rennie and Buchanan-Smith (2006) posit that it is the selection, training, and communication amongst staff that needs to be targeted in order to reduce negative and promote positive effects of humans on captive non-human primates. Potential staff must have an interest in and respect for research animals and scientific protocols and adequate training must ensure that they understand both the animals’ behavior and all protocols in which they will participate (Rennie and Buchanan-Smith 2006). It is under these conditions that a positive bond may form between humans and the individuals in their care (Rennie and Buchanan-Smith 2006). As humans learn to recognize and understand individual primates, the primates in their care do the same. Though few studies have explored this idea in depth, chimpanzees have been shown to have preferences for humans with which they have had positive interactions (Bloomsmith et al. 1997) and stump-tailed macaques have shown aggressive and fearful behaviors in response to humans with which they have had negative experiences (McKinley 2004). Proper non-human primate care should be rooted in the relationships which form between each human and each individual non-human primate (Waitt et al. 2002). Two ways of providing this are through the use of unstructured human interaction and the use of species-specific behaviors.

Unstructured human interaction describes informal “hanging out” sessions between humans and captive primates through cage barriers. It consists of unstructured bouts of play, stroking and scratching, and talking, chosen in response to the primate’s reaction (Manciocco et al 2009). Often combined with treat provisioning (Baker 2004, Baker and Springer 2006, Chelluri et al 2013, Nishimoto et al. 2015), the goal of unstructured human interactions is to utilize organic relationships that form between humans and captive primates to improve welfare, without the use of formalized training. Unstructured human interaction combined with treat provisioning has been found to improve affinity to humans, leading primates to be more likely to accept treats from familiar (Nishimoto et al. 2015) and unfamiliar humans (Baker and Springer 2006). In addition to the benefits unstructured human interaction may have on a monkey’s immediate response to humans, this form of enrichment has been found to have positive effects on primate behavior and interactions with both humans and conspecifics. Both chimpanzees and gorillas have been showed to display lower levels of abnormal behaviour in response to this type of enrichment, while increasing positive interactions with human caregivers (and with conspecifics in the case of the chimpanzees) (Baker 2004, Chelluri et al. 2015), The only study to look at the effect of unstructured human interaction without the addition of treats was completed on common marmosets by Manciocco et al. (2009). Results showed that the treatment provided led to lower levels of locomotion and self-scratching and increased levels of grooming and play behaviors amongst families when no humans were present (Manciocco et al. 2009). Unstructured human interaction therefore has the potential to greatly improve primates’ relationships with both humans and conspecifics.

The use of species-specific behavior in human interactions with captive primates is very similar to unstructured human interactions, with the key difference that primate behaviors, rather than human ones, are utilized. Treatments that utilize species-specific behaviors attempt to incorporate specific primate behaviors and contexts during interactions, taking cues from primate reactions. Behaviors are species-specific, but usually incorporate grooming, affiliative, submissive, and/or play behaviors (Bayne et al. 1993, Jensvold 2008, Jensvold et al. 2010). The theory behind using species-specific behavior is based on the idea that when dealing with primates, we are dealing with wild animals with behavioral repertoires vastly different from our own. For example, while a smile might be a very friendly behavior to a human, grins that display one’s upper teeth are considered threatening to the majority of primates (Jensvold 2008, NC3Rs “Expressions”). By using species-specific behavior one can therefore mitigate possible adverse effects of species miscommunication and attempt to meet the primates on their level, thus facilitating positive relationships.

Very few studies have looked at the immediate responses of primates to species-specific interactions with familiar humans during treatment application itself, and those that have contain somewhat ambiguous results for its effectiveness. When comparing human behavior to caregiver use of chimpanzee behaviors (posture, facial expressions, and vocalizations), Jensvold (2008) and Jensvold et al. (2010) found that chimpanzees both distinguish between and respond differently to each treatment, though neither paper concluded one was definitively preferred. Looking to longer-term effects of using species-specific behavior on the welfare of non-human primates, few studies exist that highlight this point alone, and those that do often combine the use of species-specific behavior with other treatments, such as treat provisioning or Positive Reinforcement Training (PRT). In one of the earliest explorations of species-specific behavior, combining submissive behaviors with treat provisioning, singly-housed adult Rhesus macaques saw a reduction in both frequency and duration of negative self-directed and stereotypic behavior (as well as abnormal behaviors more generally) and an increase in self-grooming behavior. In addition, many of these effects continued to be shown even after treatment ended, implying that it had long-term positive effects on the monkeys (Bayne et al 1993). When species-specific behaviors (this time gorilla play) were combined instead with Positive Reinforcement Training (PRT), they led to decreased levels of abnormal and agonistic behaviors, as well as increased levels of feeding behaviors, individual and social behaviors, affiliative behaviors, and, most notably, social play in lowland gorillas over the course of the experiment (Carrasco et al. 2009). Existing studies thus indicate that the use of species-specific behavior by care personnel has the potential to show marked behavioral improvements for many captive primates.

The main goal of this paper is to evaluate the effects of human interaction as a source of social enrichment for pair-housed female Rhesus and Cynomolgus macaques, and to see whether the monkeys respond better to human interaction, where caregivers use human behaviors during interactions, or monkey-like interaction, where caregivers attempt to incorporate monkey behaviors during interactions. This can be broken down into two main objectives: 1) Evaluate the short-term effects of both treatments during interactions, and establish monkey preference for each treatment. 2) Evaluate the long-term effects of each treatment on activity budget and presence of abnormal behaviors when no humans are present (outside of treatment application time).

## 2. Material & Methods

### 2.1 Ethics Statement

The use of animals and all procedures were approved by the Animal Care Committee of McGill University and affiliated hospitals and research institutes (protocol #2020-8155).

### 2.2 Study Population

Sixteen adult Rhesus or Cynomolgus Macaques, aged between 6 and 15 years were selected to be enrolled in the study. All monkeys were housed in Continuous Full Contact pairs in either two cages (2.03m X 1.14m X 0.91m) side-by-side or one cage and one exercise enclosure (2.49m X 1.79m X 2.11m). Monkeys were split between two rooms, and cages were distributed around the room so that all monkeys were in visual/auditory contact, but separate pairs had no physical contact. All cages were fitted with structural and occupational enrichment (i.e wooden perches, hammocks, puzzle feeders, mirrors, etc…) that were subject to change at each full cage cleaning. Full cage cleanings were done every week, during which monkeys were removed from cages, cages were disinfected, and all enrichment/floor substrate was replaced, while smaller cleanings were completed every few days to replace substrate and ensure the general cleanliness of the environment.

Artificial lighting was turned on at 7:00AM and off at 7:00PM. Monkeys were provided with a daily ration of monkey chow in the morning. Additional fruit and vegetables were provided through the caging twice daily, once at 8:00AM during positive reinforcement training with the veterinarian technicians, and once at 1:00PM by 1 of 5 caregivers, each with 1-12 years experience working with the macaques and training in basic macaque behavior. All staff typically ended their workdays by 3:00PM, at which point the monkeys would remain largely without human contact until the next day.

Of the 16 Macaques, many were already enrolled in existing biomedical studies. One to two times weekly, monkeys would be removed from their home cages and brought to a separate lab for a few hours to take recordings. Monkeys on different trials missed 1-5 days of treatment application, though never on two consecutive days. We did not consider this to be an issue for our study, as the goal was to study the effects of the treatment on laboratory-housed macaques. In order for the treatment to be truly effective, it must apply to monkeys under regular laboratory conditions, which includes being part of existing projects.

### 2.3 Data Collection

The study was conducted between August 12 and October 30, 2020, at the Comparative Medicine and Animal Resources Center (CMARC) of McGill University (Montreal, Quebec, Canada). Sixteen adult Rhesus or Cynomolgus Macaques were enrolled in a nine-week experiment (plus 2.5 week habituation) consisting of three equal phases: baseline (PRE), treatment application (EXP), and post-experiment (POST). Of the sixteen macaques, 8 adult females were selected as focal monkeys for the study, while the monkeys they lived with were treated as companions to the focal monkeys. The 8 adult females selected as focal monkeys were blocked by species and randomly assigned to one of the following treatments: Human behavior (HB) or Monkey-like behavior. The trial first began with a thirteen-day habituation (2.5 wks), in which monkeys were familiarized with all experimental procedures and materials, such as camera set-ups, coroplast dividers, etc…. Treatment was only applied during the EXP phase, while PRE and POST represented baseline and post-experimental condition, respectively. For the purpose of this project, we adapted the usual 1:00 PM feeding time of Monday through Friday to fit our experimental design, while maintaining the general daily routine the monkeys were accustomed to. No treatment occurred on weekends, which already had different schedules. Caregivers wore their regular PPE, with a few notable exceptions. Instead of their usual N95 masks and visors, caregivers wore full-face shields that filtered air and allowed their full faces to be visible. Fake-fur wrist bands were also worn to allow for grooming with the monkeys without actual contact.

Normal feeding procedures were modified to prevent monkeys from adjacent cages from seeing treatment application. A coroplast divider was introduced, and monkeys were fed one at a time, starting with the leftmost cage. During the PRE and POST phases, the caregiver and researcher would enter the divider and proceed to feed the monkey as they usually would on any normal day, for approximately 2 minutes. During the EXP phase, the caregivers followed an almost identical routine, with the addition of treatment application.

The treatment application consisted of: once in front of a cage, the researcher would start a timer and the caregiver would follow three food-times (6 minutes total): they would spend two minutes attempting to interact with the monkey with no food, then two minutes interacting while handing out produce, and finally another two minutes attempting to interact without food being provided. This was done both to see if there was an effect of the food itself versus the treatment and to see if there was an optimal time to provide social enrichment. The type of interaction provided depended on the treatment each monkey was assigned to. The goal of this design was to see both the effects of each treatment as well as the effect of extra interaction time provided to the monkeys during the EXP phase.

For both treatments, the goal was to encourage unstructured, monkey-human interactions. For monkeys receiving HB, caregivers would use only human behaviors while interacting with the monkeys. Talking calmly and kindly to the monkeys was encouraged. For monkeys receiving MB, caregivers would imitate monkey behaviors in their interactions. Talking and direct eye-contact was strongly discouraged unless absolutely necessary. All caregivers received training prior to the beginning of the trial to familiarize themselves with each treatment and all experiment procedures. Caregivers were taught three behavioral contexts to utilize: affinitive social/greet, grooming, and submissive/uninteractive (see Supp. 1.). Caregivers would begin with an affinitive social/greeting context, then they would pick which context to use based on the monkey’s context. If the monkey acted aggressive, caregivers would utilize a submissive/uninteractive behavioral context. If the monkey elicited grooming, either by offering a body part of approaching and touching the caregiver, the caregiver would use the grooming behavioral context. Finally, if the monkey acted friendly/neutral, the caregiver would first use the affinitive social context for approximately 30 seconds, then attempt to encourage more interaction by switching to a grooming context (MB).

### 2.4 Observation Procedures

A comprehensive ethogram was created based on Altman 1962, Bayne et al 1991, Partan 2002, Baker et al 2009, Kemp 2014, Cannon et al. 2016, Stolar 2018, and the NC3Rs website on Macaque care and behavior. Pretests during the habituation and pre-trial videos led to the final ethogram Table 1 and Table 2 used for this trial (for a more detailed explanation of ethogram creation, see supplemental material).

**Table 1.** Ethogram for Live Observations.

| Behavioral Categories | Behavior | Description |
|---|---|---|
| Inactive | Neutral | Monkey not engaged in any clear activity or paying attention to caregiver |
| Active |  |  |
| Eating | Eating | Feeding behaviors, including eating, foraging, and drinking. Common usage, was not coded for if monkey appeared to be chewing at beginning of video |
|  | Beg | Monkey extends hand to request food or other object from caregiver (observed in feeding context) |
| Locomotion | Locomotion | Walk, climb, jump |
| Self-directed |  |  |
| Grooming | Scratch | Repeated vigorous strokes of the hair |
|  | Self-groom | Picking, stroking, and/or licking of one's own body hair |
| Abnormal | Abnormal | Qualitatively species-inappropriate behaviors, including stereotypic, appetitive, non-injurious self-directed and self-injurious behaviours |
| Social/Interactive |  |  |
| Human Int | Interaction with Caregiver | Monkey is engaged/interacting with the caregiver |
| Monkey Int | Positive |  |
|  | Interacting with monkey positively | Affiliative behaviors directed at another monkey |
|  | Negative |  |
|  | Interacting with monkey negatively | Aggressive or submissive behaviors directed at another monkey |
| Other | Manipulate object | Using hands, feet or mouth to explore inanimate objects or caging |
|  | Not visible | Counted when the monkey was covered enough such that its behavior was not visible (ex. The head could be seen popping up behind a barrier but the body language/activity was not visible to the observer) |

**Table 2.** Ethogram for Video Observations.

| Behavioral Categories | Behavior | Description |
|---|---|---|
| Abnormal |  |  |
| Cage-directed | Lick non-food item | Licking or biting surfaces of environment, walls, or cage |
| Self-directed |  |  |
|  | Pluck Hair | Pluck hair from self, using hands or teeth |
|  | Self-bite | Any vigorous biting of one's own body |
|  | Self-clasp | Clutching or grasping own body |
| Stereotypic | Abnormal Posture | Hold seemingly uncomfortable or unnatural posture |
| Locomotive | Abnormal Locomotion | Circle, pace, rock, swing, somersault, spin, flip |
|  | Pacing | Repetitively walking or running back and forth in a circle |
|  | Rocking | Any rhythmic motions of the body from a stationary position; animal remains sitting or standing while the upper torso sways back and forth |
| Social |  |  |
| Positive |  |  |
| Affiliative | Follow | Walk close behind another monkey, same direction |
|  | Lipsmack | Rapid lip movements, with or without tongue, directed towards a conspecific or person |
|  | Play | Complex form of behavior interactions involving many of the other behavior patterns in highly modified form. (social or solitary) |
|  | Playface | The ears and brow are pulled back, the mouth is open and the top lid is pulled over the teeth |
|  | Present | Present part of body (neck, side, or often backside) to caregiver, looking away and becoming still. Often in grooming context, where animal will approach or reposition itself in front of another, presenting an area of the body to be groomed. Also a sign of submission or sexual behavior |
|  | Touch | Monkey has physical contact with another individual or caregiver. Not aggressive |
| Grooming | Allogroom | Hands systematically combing through other monkey's hair (One animal picks through the hair of another with hands or teeth, removing skin, dirt or ticks) |
|  | Self-groom | Picking, stroking, and/or licking of one's own body hair |
| Negative |  |  |
| Aggressive | Bite | Focal individual puts mouth and teeth on body part or clothing of an individual |
|  | Branch Shake | Bounce up and down on branch or other object, holding on with feet so that the substrate moves as well |
|  | Chase | Run after other monkey (dominant animals will chase lower ranked animals away from resources where there is high competition, and use aggression to gain access.) |
|  | Displace | Focus animal moves to where another is sitting who then moves away (or opposite) |
|  | Fight | Fast-paced agonistic interaction involving chasing, tumbling, biting, barking, screaming all at once |
|  | Grab | Use hand to grab hold of other monkey |
|  | Head bob (and thorax) | The head was jerked very quickly up and down. Arms are sometimes flexed and extended (like a push-up) |
|  | Head Jerk | Head moved quickly and abruptly up and down |
|  | Hit | Meaning obvious/use hand/arm to hit another monkey |
|  | Lunge | Sudden and quick movement of upper or entire body towards recipient |
|  | Open Mouth Threat | Almost always accompanied by direct stare, often by head bobbing. The mandible is opened, but the lips are not bared. The tips of the canine of an adult male are visible but not conspicuous during this gesture. (The open mouth stare is used to threaten other individuals. It is seen during tension between group members, and is often performed towards unwelcome humans) |
|  | Push | Meaning obvious, use hand/arm to push another monkey |
| Other |  |  |
| Self-grooming | Self-groom | Picking, stroking, and/or licking of one's own body hair |

Scans of general activity were taken every 15 seconds on focal monkeys during the six minutes of treatment application for each monkey daily. Data was collected by one observer. One-hour-long videos using GoPro (HERO Black 7) on tripods were recorded every Friday afternoon from 4:00 PM to 5:00 PM across all three phases. The goal was to observe any long-term effects of either or both treatments on monkey social and abnormal behavior when no humans were present.

### 2.5 Data Analysis

#### 2.5.1 Live Observations

Weekly averages were calculated for each variable to control for the effect of caregiver.

This was done because while 5 different caregivers participated in the study, they were distributed across each week such that while individual days were influenced by caregivers, whole weeks were identical in the distribution of caregivers. Measurements are presented as proportion of time (duration in sec divided by total observation time, i.e. 6 min or 360 sec).

#### 2.5.2 Video Recordings

Video recordings were analyzed by one trained observer. The observer was trained and assessed on reliability from video recordings of pre-trial data. Overall inter- and intra-reliability of at least 80% was obtained using kappa coefficients once before scoring of trial data commenced and was retested twice throughout scoring (approximately every 2-3 weeks). The observer recorded the start and end time whenever the focal monkey used one of the six behavioral contexts of interest: Abnormal Behavior, Affiliative Behavior, Grooming Behavior, Agonistic Behavior, Submissive Behavior, and Self-Grooming Behavior (see Video -Table2). Other behavioural contexts were not recorded. Continuous observation of each behavioral context was completed for the full hour. A behavioral context was considered to begin when a behavior unique to that context began. A context was considered to end or change when either no behaviors from that context were being performed or when a behavior from a different context was spotted, at which point that context would begin.

Frequency and duration were extracted from data. In addition, durations of behavioral contexts were divided by duration of videos, in all cases one hour, or 3600 seconds in order to produce a proportion of time spent in each context.

#### 2.5.3 General Model Considerations

Given that only 8 monkeys were enrolled in the study to split between the two treatments, where 5 were Rhesus Macaques and the remaining 3 were Cynomolgus, it was not possible to analyze the effect of species. Instead, species was treated as a block effect for analyses. This allowed us to account for the effect of species while still focussing on our main question: assessing the effect of treatment. Only weekly averages were used in the final results to eliminate the need to include caregiver in the final models. Significance was set at 0.10. All models were run using SAS 9.4 and tested using PROC MIXED and PROC UNIVARIATE procedures. Results are presented as means with standard error.

### 2.6 Statistical Models

The data that support the findings of this study are openly available in GitHub at https://github.com/RachelvanVliet/NHP-Project.git (van Vliet et al. 2020).

#### 2.6.1 Live Observations

For live observation analysis, food-time (accounting for the 6 minutes of interactions around which food is provided, specifically before, during, and after food distribution) and week were considered independent of one another. Since we were explicitly interested in the effect of food-time over and above the effect of week, both are included in the model as main effects. In order to account for their relationship and place in context, a random interaction between food-time and week was included. Food-time, as a variable of interest, was included as a fixed effect, while week was considered random. Monkey was considered a random effect as we are not interested in the individual monkeys themselves and they represent significant variation in the model.

The following model was therefore developed to analyze the live observations:

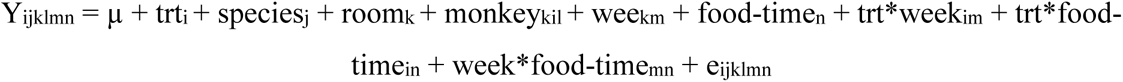

Where: Y_ijklmn_ is the dependent variable, which is the outcome measure of the l^th^ monkey from the j^th^ species and k^th^ room, receiving the j^th^ treatment at the n^th^ time during the m^th^ week. The trt_i_ is the fixed effect of the i^th^ treatment (j = HB or MB). The species_j_ is the fixed (block) effect of the j^th^ species (j= Rhesus or Cynomolgus macaque). The room_k_ is the fixed effect of the k^th^ room (k=1421B or 1421E). The monkey_kil_ is the random effect of the l^th^ monkey (l= 1 to 8) from the j^th^ species and k^th^ room. The week_m_ is the random effect of the m^th^ week (m= 1, 2, or 3). The food-time_n_ is the fixed effect of the n^th^ food-time (n= before, during, or after feeding). The trt*week_im_ is the random effect of the interaction between the i^th^ treatment and the m^th^ week. The trt*food-time_in_ is the fixed effect of the interaction between the i^th^ treatment and the n^th^ food-time. The week*food-time_mn_ is the random effect of the interaction between the m^th^ week and the nth food-time. Finally, e_ijklmn_ is the random error.

In order to account for repeated measures of monkey and week, the best fitting covariance structure (between Auto-Regressive (1) or AR(1) and Compound Symmetry or CS) was selected with a step-by-step elimination process of inclusion and exclusion of different random and repeated factor combinations. The chosen combination for each outcome measure was decided by the best BIC fit statistics. To ensure that the data came from a normal population, for each outcome measure, normality of residuals for each outcome measure was tested using PROC MIXED and PROC UNIVARIATE procedures. A Tukey adjustment was used in order to account for the multiple comparisons associated with each interaction.

#### 2.6.2 Video Recordings

For weekly videos, week was considered nested in phase, as the effect of week is only of interest, and in fact only comprehensible, when nested within each phase. As for the live observation analysis, room was considered as a fixed effect and monkey as a random effect. No interaction was included between treatment and week. Week is not of particular interest to the research question (unlike phase), and upon running the first few variables with the interaction where none was found to be significant, it was removed from the model.

In order to analyze the weekly videos, the following model was developed:

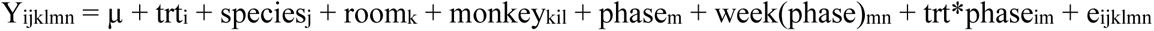

Where: Y_ijklmn_ is the dependent variable, which is the outcome measure of the l^th^ monkey from the j^th^ species and k^th^ room, receiving the i^th^ treatment during the nth week of the m^th^ phase. The trt_i_ is the fixed effect of the i^th^ treatment (i = HB or MB). The species_j_ is the fixed (block) effect of the j^th^ species (j= Rhesus or Cynomolgus macaque). The room_k_ is the fixed effect of the k^th^ room (k= 1421B or 1421E). The monkey_kil_ is the random effect of the l^th^ monkey (l= 1 to 8) from the j^th^ species and kth room. The phase_m_ is the fixed effect of the m^th^ phase (m= PRE, EXP, and POST). The week(phase)_mn_ is the fixed effect of the n^th^ week (n = 1, 2, or 3) from the m^th^ phase. The trt*phase_im_ is the fixed effect of the interaction between the j^th^ treatment and m^th^ phase.. Finally, the e_ijklmn_ is the random error.

To account for repeated measures, the best fitting covariance structure (between Auto-Regressive (1) or AR(1), Compound Symmetry or CS, and Unstructured or UN) was selected for each outcome measure by comparing the BIC fit statistics. Normality of residuals for each outcome measure was testing using PROC MIXED and PROC UNIVARIATE procedures. A Tukey adjustment was used in order to account for the multiple comparisons associated with each interaction.

## 3. Results

### 3.1 Live Observations

Few differences were found between treatment groups on general behavior scans taken during treatment. An effect of treatment was found for abnormal behaviors, in which MB monkeys spent 1.0 ± 0.23% (or 3.42 sec, F22.9 = 16.61, p = 0.0005) less proportion of time performing abnormal behaviors than HB monkeys (for a full summary of results, see Supp. 2).

There was a consistent effect of food-time found for most outcome measures. For monkeys from both groups across all weeks, inactive behavior was highest during the first two minutes of observation (before feeding) and lowest during the second two while food was being handed out. Conversely, active behaviors were lowest in the first two minutes and highest during the second. Within active behaviors, this trend holds true only for eating, which is the main outcome measure causing this tendency. Monkeys spent 82.0 ± 1.50% (F40 = 607.15, p < 0.0001) of their time eating while food was being provided, which was higher than both before and after, though monkeys from both groups still spent 60.4 ± 1.50% (F40 = 607.15, p < 0.0001) of their time continuing to eat after food was no longer provided. For other active behaviors, the opposite trend was observed. Monkeys had higher rates of locomotion, self-directed behaviors, and social/interactive behaviors in the first two minutes of interaction, before food was provided (see Supp. 2).

For self-directed behaviors, there was also an effect of week, where monkeys showed a 1.1 ± 0.47% increase in self-directed behaviors from week one to week three (F11.2 = 3.68, p = 0.081).

Within self-directed behaviors, there was an effect of the food-time*week interaction for self-grooming. Monkeys still spent the most time displaying self-grooming behaviors before food was provided, but in addition self-grooming increased by 2.2 ± 0.69% (7.9 sec, F52 = 2.04, p= 0.058) during this time from week one to three. Within social/interactive behaviors, effects of food-time were also found. For time spent interacting with caregivers (Human-int), a numerical difference indicates that monkeys from both groups once again showed higher rates before feeding than after and lowest rates while food was provided, though upon further analysis no two food-times were found to be significantly different. For Monkey-int, or time spent interacting with conspecifics, an effect of food-time was found, though no two food-times were found to be significantly different from each other. Within monkey interaction behaviors, however, an effect of the food-time*week interaction was found for negative interactions. During week 1 of treatment application these were highest in the first two minutes of interaction than either during food provisioning (by 3.1 ± 0.10%, F40 = 3.13, p = 0.068) or after (by 3.4 ± 0.10%, F40 = 3.13, p = 0.031). In addition, negative monkey interactions before feeding saw a 3.4 ± 1.13% (or 12.4 sec, F40 = 3.13, p = 0.087) decrease in negative interactions over the course of the three weeks of treatment application.

### 3.2 Video Recordings

Amongst the three major categories observed for weekly videos (abnormal, social, other), no effects were found for frequency, and no effects were found for proportion of time for either abnormal or other behavior. For social behavior, an effect of phase was found for the proportion of time spent in this context. Monkeys across both treatments saw a 12.6 ± 6.10% (7.56 min, F54 = 2.41, p = 0.107) increase in proportion of time performing social behaviors from phase one to three.

When looking at the distribution of behaviors within the major categories, effects were found for both abnormal and social behaviors. Within abnormal behaviors, effects of both treatment and phase, though not their interaction, were found for cage-directed behaviors. Overall, MB monkeys spent a lower proportion of time performing cage-directed behaviors than HB monkeys (lsmeans MB: 2.0 ± 0.44 %vs HB: 3.4 ± 0.44%; F4 = 4.71, p = 0.0957).

However, monkeys from both treatments saw a decrease of 3.5 ± 1.49 (F54 = 3.34, p = 0.0584) in the frequency of cage-directed behaviors performed across the three phases. There was also an effect of phase on the frequency of stereotypic behaviors performed. Monkeys from both treatments performed 1.2 ± 0.53 (F54 = 2.78, p = 0.0668) fewer bouts of stereotypic behaviors from phase two to three. No further effects were found for either self-directed or locomotive behaviors.

Within social behaviors, several effects were found. While overall social behaviors increased across all three phases for both treatments, this trend was only true for positive social behaviors, for which there was an effect of phase on both frequency and proportion of time. For frequency of positive behaviors, a significant increase of 3.0 ± 1.43 between phases two and three was found (F35.1 = 2.54, p = 0.0997, see table 3.2.4.). Monkeys from both treatments further saw a 12.9 ± 6.12% (or 7.71 min, F54 = 2.40, p = 0.0996) increase in the proportion of time spent socializing positively with conspecifics from phase one to three (see table 4). A significant effect of phase was found for Negative Behaviors, though displaying the opposite trend Monkeys from both groups saw a 0.7 ± 0.30% (24.84 sec, F54 = 2.73 p = 0.0643) decrease in the proportion of time spent interacting negatively from phase two to three (see Supp. 2).

Within positive behaviors, an effect of phase was found for both frequency and proportion of time monkeys spent performing grooming behaviors. Monkeys across both treatments increased the frequency and proportion of time spent grooming across all three phases (see Supp. 3). Monkeys performed 1.6 ± 0.80 (F53.8 = 2.34, p = 0.1073) more bouts of grooming and spent 10.6 ± 4.78% (F54.1 = 2.47, p = 0.0766) more of their time grooming from phase one to three (see figure 1). No effects were found for affiliative behaviors.

**Figure 1.**
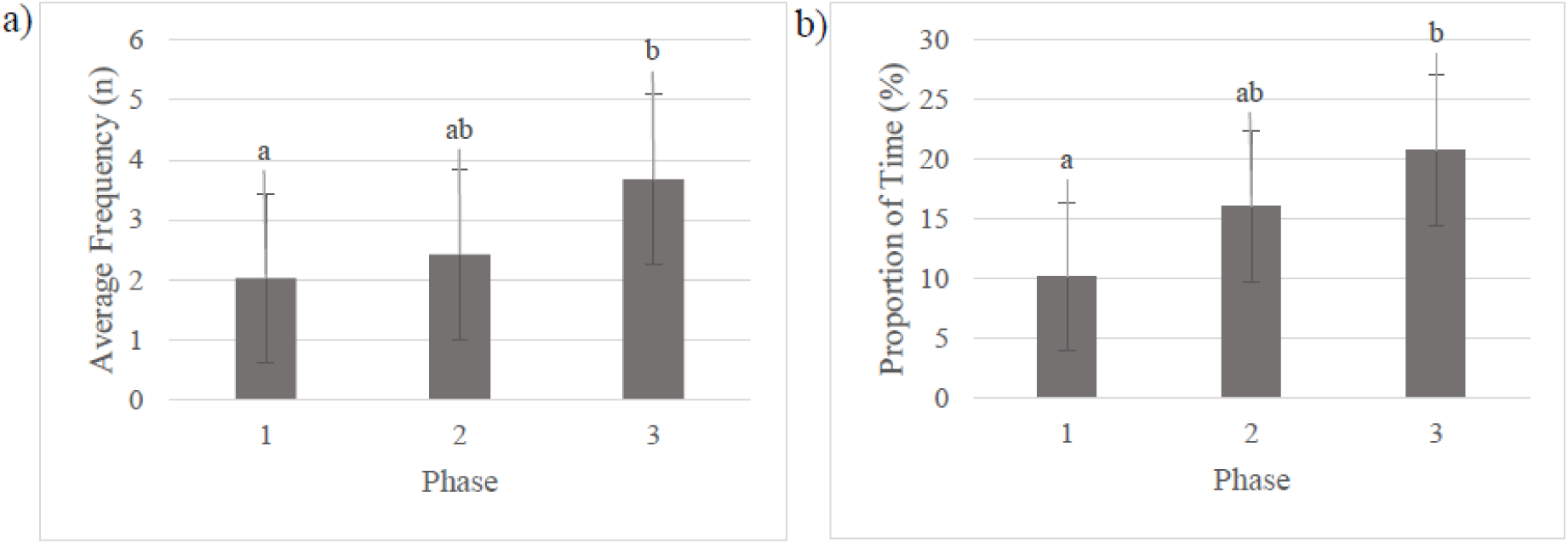
Frequency of Grooming (a) and Proportion of Time Grooming (b) across phase. Bars represent estimated means of the model and SE. Bars with different superscripts (a, b, c) differ (Post hoc Tukey test, P < 0.10).

Within negative behaviors, a similar trend to the larger group was found for aggressive behaviors. A significant decrease from phase two to three was found. Monkeys from both treatment groups saw a 0.5 ± 0.25% (19.44 sec, F54.1= 2.31, p = 0.0897) decrease in the proportion of time spent performing aggressive behaviors towards conspecifics from phase two to three. No effects were found for submissive behaviors.

## 4. Discussion

For almost every behavior looked at during scans, there was a significant effect of what we called ‘food-time’, or the distribution of time around when food was provided. With the exception of eating (which, unsurprisingly, peaked while food was distributed), all behaviors were highest in the first two minutes of interaction. Monkeys, either waiting for food to be distributed and looking for distractions, genuinely interested in the caregivers then distracted during and after feeding by food, or distracted by other things until food was provided, were most active in the first two minutes of treatment application. This finding was unsurprising, as food has been found to be one of the most engaging forms of enrichment that has the greatest influence on monkey time budgets (Jaman and Huffman 2008), which would explain the dominance of eating during (and possibly after) feeding. Many studies that have looked at human interaction as a form of enrichment or to ameliorate human-primate relationships have combined treat provisioning with interactions (Bayne et al. 1993, Baker 2004, Baker and Springer 2006, Chelluri et al. 2015, Nishimoto et al. 2016). Our results, however, imply that many possible behaviors and reactions primates may have to the interactions alone, may have been masked by the influence of the food being provided. It further implies that future research that aims to maximize the benefits of caregiver interaction should perhaps focus on the time before feeding, rather than during or after.

One of the most notable results in our study is the impact both treatments had on grooming, both during the live scans and in the weekly videos. There was a noted increase in self-grooming behavior before food provision by caregivers during treatment across the three weeks of treatment application. Moreover, in weekly videos (after treatment application with no humans present), the proportion of time performing social behaviours increased. Within this, it was positive behaviours, and, most notably, allogrooming that saw an increase in both frequency and proportion of time across all three phases.

One of our main hypothesis is providing additional social contacts to the monkeys would result in increased positive engagements between monkeys (as well as reduced abnormal behaviors). The main treatment we applied was, ultimately, grooming. Caregivers did not have access to other forms of physical touch and therefore positive interactions such as play were not possible. Instead, the main positive interaction provided was either grooming or offering/being groomed. Increases in allogrooming after the application of human interaction has been also observed in common marmosets (Manciocco et al. 2009) and chimpanzees (Baker 2004). It is interesting to note, however, that when species-specific play behaviors (rather than grooming) were administered as the main treatment to captive lowland gorillas, Carrasco et al. (2009) found that levels of both individual and social play increased. In this case, therefore, any increase in grooming behavior, whether social or not, implies that the treatments were having an impact on monkey behavior by encouraging them to spend more of their time grooming as opposed to other activities. Moreover, grooming is a highly important behavior for macaques (Jaman and Huffman 2013). While social grooming is used to strengthen bonds and group cohesion (Jaman and Huffman 2013), self-grooming may have both self-cleaning and self-calming purposes (particularly in times of excitement) (Bayne et al. 1993).

This argument is further strengthened when we look at the opposite trend for negative interactions during live scans: monkeys were replacing negative interactions with conspecifics with self-grooming during the first two minutes of treatment application across the three weeks. The positive social interactions with caregivers may have, over time, encouraged more friendly relationships between monkeys, resulting in less confrontation when a defendable resource (either the caregiver or food) is provided. A similar tendency has been found for lowland gorillas, in which the addition of Positive Reinforcement Training (PRT) and ‘gorilla play’ resulted in an overall decrease in agonistic behavior between apes (Carrasco et al. 2009). It is equally possible that this decrease represents instead a slow habituation to the treatment and the new schedule. Indeed, monkeys were initially used to being fed immediately. The initial change to two minutes without food may have caused some frustration, fear, or competition between conspecifics, resulting in increased negative interactions, as has been seen in stump-tailed macaques when feeding times become inconsistent or delayed (Waitt and Buchanan-Smith 2001). As the monkeys adapted to the new routine, however, this fear/frustration, and as a result the negative interactions, waned. It could also be a combination of the two, where monkeys both adapted to later feeding and the extra time with caregivers encouraged less negative interactions.

During weekly videos, positive social interactions between monkeys from both treatments saw an increase in the proportion of time spent socializing positively across all three periods, as well as an increase in the frequency of these positive interactions between phases 2 and 3, supporting the idea that both treatments led to an overall increase in positive social interactions. Conversely, monkeys from both groups showed a decrease in the proportion of time spent interacting negatively, and within negative behaviors the same can be said for aggressive behaviors, between phases 2 and 3, providing further evidence that the positive benefits exceed any potential negative effects of both treatments.

Another major finding was the impact each treatment had on the expression of abnormal behaviours, which were lower amongst MB monkeys than HB monkeys in both live observations and videos, and decreased in videos over the course of the experiment. These videos were taken in the afternoon at a time when no humans were present, and therefore serve to provide an indication of the long-term effects of treatment application, beyond when it is actually provided. Overall, MB monkeys showed lower levels of cage-directed abnormal behaviours than HB monkeys. Abnormal behaviours develop in response to either aversive environments or those lacking stimulation (Mason 1991). Their continued presence or change in their occurrence in response to a treatment, is therefore an important indicator of welfare in conjunction with other findings. Their lower frequency amongst MB monkeys thus indicates a possible benefit of this treatment. We further found an effect of phase, in that regardless of treatment, all monkeys saw a reduction in cage-directed behavior across all three phases. A similar trend was found for stereotypic behavior. This is a very positive result, as a reduction in abnormal behaviors is one of the major welfare measures used for lab animals (Prescott et al. 2022).

Unlike Bayne et al. (1993)’s study providing a similar treatment to individually-housed rhesus macaques, we did not find more pronounced effects on all types of abnormal behaviors. While this initially may seem to be a disappointing result, it is important to note the differences between our study populations. Bayne et al.’s monkeys were individually housed, received no other enrichment, and had high baseline levels of the abnormal behaviors being targeted. The monkeys present in our study, by contrast, were already in social housing (pairs), received enrichment daily, and showed low baseline levels of each type of abnormal behavior. It is therefore likely that the monkeys enrolled in this study simply did not have high enough baseline levels of abnormal behavior for significant changes to occur only with the treatment applied. It is therefore even more interesting that both cage-directed and stereotypic behaviors were still found to be affected by the extra time provided by both treatments. This idea is further strengthened by a study done by Baker et al. (2009) using PRT on macaques, in which noted reductions in rates of abnormal behaviors were only seen for monkeys that had a high level to begin with. This does not mean that the enrichment was not beneficial, however, but simply that it alone would not be enough to show significant changes in these types of behaviors, at least for the amount of time it was applied. Alternatively, it may be the case that the type of treatment applied simply did not affect all types of abnormal behaviors, and future work should focus more specifically on cage-directed and stereotypic behavior. An interesting question for future work would look to our treatments in conjunction with other interventions or enrichments or applied for longer periods of time in order to see if caregiver interaction can contribute to larger reductions in abnormal behaviors.

## 5. Conclusion

Our project sought to evaluate the use of caregivers, who work with and are in regular contact with the macaques in their care, as a form of social enrichment. During the three weeks of treatment application, where the macaques received 6 additional minutes of either human or monkey-like interaction, macaques engaged most with caregivers in the 2 minutes before food was provided, rather than during or after. While many studies have combined human interaction with food provisioning, our results indicated that the most effective time to engage is in fact before food is provided. Over the course of the three weeks, monkeys from both treatments saw an increase in self-grooming behavior and, conversely, a decrease in negative interactions with other monkeys. Over the course of the whole experiment (9 weeks), we once again noted an increase in grooming behaviors, though this time between conspecifics. In addition, positive social interactions increased while negative interactions and cage-directed negative behaviors decreased. Between treatments, MB monkeys displayed lower rates of abnormal behavior during interactions and lower rates of cage-directed behaviour in weekly videos.

These results suggest that macaques in laboratory environments benefit from social interaction with caregivers, though we did not conclusively find that either human or monkey-like behavior was more adapted. Though MB monkeys displayed slightly lower rates of abnormal behavior in response to treatment, very few other treatment differences were found. Future studies should look to increasing treatment and experiment time in order to see if more time would create more pronounced results. Our study points to future directions for management and care practices in laboratory environments for rhesus and cynomolgus macaques. It further implies that humans can be effectively used to provide social enrichment for captive primates within the constraints of laboratory environments.

## Acknowledgements

We would also like to thank Dr. Roger Cue and Tania Wolfe for their help with experimental design and statistical analysis elaboration. Thanks to Simran Prassad for her help with elaborating the ethogram, data collection and handling, and video watching. We are also thankful to the Comparative medicine and Animal Resources Center (CMARC) staff, Lori Ann Burgess, Gianfranco Galardo, Philippe Laroche, Christopher O’Hare Friere, Jeremie G. Smith, Josef Penalosa Nacario, Mitchell Joseph McKinnon, and Pacholo Infante Libarnes who contributed a great deal to the success of the project. We would also like to thank Jim Gourdon for all of his help with the elaboration of the study and funding acquisition.

