## Supplemental Materials for "Making monkeys of caregivers: Evaluating the use of caregiver behaviour as social enrichment for lab-housed Rhesus and Cynomolgus Macaques"

Supporting Information

Github (SAS code):
https://github.com/RachelvanVliet/NHP-Project/blob/main/README.md

### Ethogram Creation

In order to create the ethogram, we began by first creating a comprehensive list of all behaviors that may be identified or seen. This was done by compiling ethograms found in Altman 1962, Bayne et al 1991, Partan 2002, Baker et al 2009, Kemp 2014, Cannon et al. 2016, Stolar 2018, and the NC3Rs website on Macaque care and behavior. Duplicate behaviors were either deleted or definitions were combined to provide the most information possible. For the live observation ethogram, the observer trained during the habituation using the comprehensive ethogram to see what behaviors could be identified during the 6-min interactions. For the weekly video ethogram, two observers trained on pre-trial videos using the comprehensive ethogram to see what behaviors would be visible/possible to code during weekly videos. When a behavioral context was seen to begin, individual behaviors seen would be listed beside the start and end time for that context. Contexts were originally sorted as abnormal, grooming, or social. They were later recoded into abnormal, social (positive-affiliative and grooming and negative-agonistic and submissive), and other (self-grooming). This resulted in the final ethogram used throughout the experiment duration.

**Supp. 1.** Caregiver Guidelines, providing behaviors to be performed in each context (chosen based on monkey's reaction).

| | HB | MB |
|---|---|---|
| Affinitive Social/Greet | Say hello, smile, talk calmly and softly to monkey, greet (say hello), laugh | lipsmacking, playface, neutral face, grunt |
| Grooming | Inspect/comb through monkey’s fur, scratch, eye contact, speak softly | Offer wrist, comb through/inspect monkey’s fur, lipsmacking |
| Submissive/Uninteractive | Maintain posture, but cower slightly, momentarily look away, continuously talk to monkey to calm down (speak slowly/soothingly) | Crouch/get low, avoid eye contact, avoid making sound, backup slightly (a step or two) |

**Supp. 2.** Behaviour Scans of Active (eating, locomotion, self-directed (self-grooming and abnormal), Social/Interactive (Human and monkey)) and Inactive behaviours, differences in proportion of time averaged across weeks between food-times.

| | Food-time Lsmeans (SE)[1] | | | Food-time | | |
|---|---|---|---|---|---|---|
| | Before | During | After | F[2] | Ddf | *P* |
| Active | 42.3 | 91.7 | 81.1 | 89.56 | 52 | <0.0001 |
| | (3.69)[a] | (3.69)[b] | (3.69)[c] | | | |
| Eating | 4.6 | 82 | 60.4 | 607.15 | 40 | <0.0001 |
| | (1.50)[a] | (1.50)[b] | (1.50)[c] | | | |
| Locomotion | 15.1 | 10.4 | 12.9 | 3.96 | 40 | 0.0271 |
| | (1.99)[a] | (1.99)[b] | (1.99)[ab] | | | |
| Self-directed | 2.1 | 0.35 | 0.76 | 5.1 | 32.4 | 0.0119 |
| | (0.40)[a] | (0.40)[b] | (0.40)[b] | | | |
| Self-grooming | 1.5 | 0.2 | 0.62 | 5.14 | 52 | 0.0092 |
| | (0.28)a | (0.28)b | (0.28)b | | | |
| Abnormal | 0.7 | 0.2 | 0.2 | 2.24 | 34.6 | 0.1215 |
| | (-0.21) | (-0.21) | (-0.21) | | | |
| Social/Interactive | 20.7 | -0.7 | 7.3 | 22.08 | 29.3 | <0.0001 |
| | (3.42)[a] | (3.42)[b] | (3.42)[c] | | | |
| Human Interaction | 18.9 | -1.3 | 6.1 | 19.51 | 32.2 | <0.0001 |
| | (-3.22) | (-3.22) | (-3.22) | | | |
| Monkey-interaction | 1.8 | 0.5 | 1.2 | 1.8 | 40 | 0.1784 |
| | (-0.72) | (-0.72) | (-0.72) | | | |
| M-Positive | 0.2 | 0.02 | 0.3 | 1.15 | 28.6 | 0.3306 |
| | (-0.2) | (-0.2) | (-0.2) | | | |
| M-Negative | 1.5 | 0.5 | 0.8 | 1.74 | 40 | 0.1879 |
| | (-0.48) | (-0.48) | (-0.48) | | | |
| Inactive | 57 | 8.2 | 18.8 | 93.34 | 52 | <0.0001 |
| | (3.57)a | (3.57)b | (3.57)c | | | |

[1] LSmeans of effects within a row with different superscripts (a, b, c) differ (P < 0.10).

[2] Where F represents F-value, Ddf represents denominator degrees of freedom, and *P* represents *P*-value

**Supp. 3.** Positive (affiliative, grooming) and Negative (aggressive, submissive) behavioural contexts, differences in frequency and proportion of time averaged across weeks between phase and treatment*phase

| | Phase Lsmeans (SE)[3] | | | Treatment*Phase Lsmeans (SE) | | | | | | Phase | | | Treatment*Phase | | |
|---|---|---|---|---|---|---|---|---|---|---|---|---|---|---|---|
| | | | | HB | | | MB | | | | | | | | |
| | 1 | 2 | 3 | 1 | 2 | 3 | 1 | 2 | 3 | F[4] | Ddf | *P* | F | Ddf[4] | *P* |
| Frequency | | | | | | | | | | | | | | | |
| Positive | 6.2 (2.34)ab | 5.4 (2.35)a | 8.5 (2.35)b | 6.7 (3.36) | 4.3 (3.36) | 7.6 (3.37) | 5.6 (3.14) | 6.6 (3.19) | 9.4 (3.19) | 2.54 | 35.1 | 0.0936 | 0.83 | 22.0 | 0.4483 |
| Affiliative | 4.2 (1.15) | 3.1 (1.15) | 4.9 (1.15) | 3.5 (1.62) | 1.4 (1.62) | 2.7 (1.62) | 4.9 (1.60) | 4.7 (1.60) | 7.0 (1.60) | 1.62 | 38.4 | 0.2114 | 1.20 | 20.5 | 0.3211 |
| Grooming | 2.0 (1.40)a | 2.4 (1.42)ab | 3.7 (1.42)b | 3.2 (2.05) | 2.8 (1.05) | 4.9 (2.05) | 0.9 (1.84) | 2.0 (1.90) | 2.4 (1.90) | 2.34 | 53.8 | 0.1061 | 0.66 | 53.8 | 0.521 |
| Negative | 4.6 (2.56) | 6.1 (2.58) | 4.4 (2.58) | 2.6 (3.75) | 2.3 (3.75) | 1.9 (3.75) | 6.7 (3.32) | 9.9 (3.38) | 6.9 (3.38) | 1.53 | 53.5 | 0.2265 | 1.44 | 53.5 | 0.2466 |
| Aggressive | 2.8 (1.32) | 3.5 (1.34) | 2.5 (1.34) | 1.5 (1.92) | 1.5 (1.92) | 1.4 (1.92) | 4.2 (1.74) | 5.4 (1.80) | 3.6 (1.80) | 0.70 | 53.9 | 0.4998 | 0.49 | 53.9 | 0.6176 |
| Submissive | 1.8 (1.42) | 2.7 (1.44) | 1.9 (1.44) | 1.1 (2.07) | 0.8 (2.07) | 0.5 (2.07) | 2.5 (1.87) | 4.6 (1.93) | 3.3 (1.93) | 0.60 | 53.8 | 0.5544 | 0.97 | 53.8 | 0.3868 |
| Proportion of time | | | | | | | | | | | | | | | |
| Positive | 14.9 (8.51)a | 24.8 (8.64)ab | 27.8 (8.64)b | 19.5 (12.36) | 21.7 (12.36) | 29.2 (12.36) | 10.3 (11.33) | 28.0 (11.80) | 26.4 (11.80) | 2.40 | 54.0 | 0.1002 | 0.81 | 54.0 | 0.4496 |
| Affiliative | 4.9 (2.92) | 9.0 (2.97) | 7.3 (2.97) | 3.7 (4.19) | 7.9 (4.19) | 2.5 (4.19) | 6.2 (4.00) | 10.1 (4.18) | 12.0 (4.18) | 0.90 | 54.2 | 0.4143 | 0.91 | 54.2 | 0.4097 |
| Grooming | 10.1 (6.20)ab | 16.1 (6.30)a | 20.7 (6.30)b | 15.7 (8.99) | 13.6 (8.99) | 26.5 (8.99) | 4.5 (8.29) | 18.4 (8.66) | 15.0 (8.66) | 2.47 | 54.1 | 0.0938 | 1.93 | 54.1 | 0.1551 |
| Negative | 0.5 (0.41)ab | 1.0 (0.42)a | 0.3 (0.42)b | 0.3 (0.60) | 0.2 (0.60) | 0.03 (0.60) | 0.8 (0.55) | 1.8 (0.57) | 0.6 (0.57) | 2.73 | 54.0 | 0.0745 | 2.28 | 54.0 | 0.1118 |
| Aggressive | 0.5 (0.31)ab | 0.7 (0.31)a | 0.2 (0.31)b | 0.3 (0.45)ab | 0.1 (0.45)ab | 0.02 (0.41)ab | 0.6 (0.41)ab | 1.3 (0.43)a | 0.3 (0.43)b | 2.31 | 54.1 | 0.1090 | 2.35 | 54.1 | 0.1050 |
| Submissive | 0.1 (0.12) | 0.3 (0.13) | 0.2 (0.13) | 0.1 (0.18) | 0.2 (0.18) | -0.03 (0.18) | 0.1 (0.16) | 0.4 (0.16) | 0.3 (0.18) | 0.99 | 16.3 | 0.3929 | 0.48 | 29.2 | 0.6248 |

[3] LSmeans of effects within a row with different superscripts (a, b, c) differ (P < 0.10).

[4] Where F represents F-value, Ddf represents denominator degrees of freedom, and *P* represents *P*-value
